# ENS lineage potential is not intrinsically regionalized but is modulated by PTPRZ1 signaling

**DOI:** 10.64898/2026.03.05.709942

**Authors:** Ali Kalantari, Ophir Klein, Zev J. Gartner, Faranak Fattahi

**Author notes:** Correspondence: Zev Gartner, Department of Pharmaceutical Chemistry, Chan Zuckerberg Biohub, University of California, San Francisco, 600 16th St., GH N512E, San Francisco, CA 94158, Faranak Fattahi, Department of Cellular and Molecular Pharmacology, Eli and Edythe Broad Center of Regeneration Medicine and Stem Cell Research, University of California, San Francisco, 600 16th St., GH S572F, San Francisco, CA 94158.

## Abstract

The enteric nervous system (ENS) orchestrates critical gastrointestinal functions including peristalsis, fluid exchange, and blood flow regulation, and develops from vagal neural crest (vNC) progenitors that colonize the gut. While the gut epithelium and mesenchyme exhibit pronounced anterior–posterior (A-P) transcriptional patterning and lineage diversification after mid-gestation, whether the ENS itself undergoes comparable regional embryonic transcriptional diversification has remained unclear. Here, we use multiplexed single-cell RNA sequencing and functional perturbations to dissect how the ENS is patterned between E13.5 and E18.5 within the context of a regionally specialized gut. We find that, while the epithelium and mesenchyme display strong and enduring AP-graded gene expression programs, the ENS lacks intrinsic regionalization and instead follows a predominantly temporal maturation trajectory characterized by neuronal and glial differentiation states. Integrative ligand-receptor analyses reveal that mesenchymal populations express A-P patterned microenvironmental cues that correlate with subtle, region-linked transcriptional tuning in ENS cells, despite the absence of intrinsic A-P identities. Among these signals, PTN/MDK-PTPRZ1 signaling emerges as a major spatial and temporal input to the ENS, with gradients that track both small intestinal region and developmental time. To test the relevance of PTPRZ1 signaling for human ENS development, we perturbed pluripotent stem cell-derived ENS cultures and found that modulating PTPRZ1 signaling impacts proliferative, neurogenic, and neurotransmitter-specification programs, confirming that niche-derived cues fine-tune ENS development. Together, our findings support a model in which the small intestine establishes A-P regionalization through epithelial and mesenchymal patterning, whereas the ENS maintains a relatively uniform core neuroglial program that is secondarily refined by localized microenvironmental signals. This framework highlights how extrinsic, region-specific cues, rather than intrinsic regional transcriptional codes, shape ENS maturation within the small intestine.

## INTRODUCTION

During embryogenesis, the gut tube emerges from endodermal and mesodermal precursors and subsequently develops into a regionally patterned organ with distinct morphological and functional domains along its anterior-posterior (AP) axis^1^. These specialized domains are shaped in part by the epithelium (endoderm-derived) and the surrounding mesenchyme (mesoderm-derived), each adopting unique transcriptional programs that drive tissue morphogenesis and organ-specific functions^2^. Combinatorial cellular diversity within and between these compartments generate discrete niches that influence the fate and behavior of other cell lineages during intestinal development. Notably, the small intestine harbors migrating progenitor populations derived from other anatomical locations, such as neural crest, endothelial, and myeloid cells, that integrate into the gut and contribute to its complexity^3^. Although A-P patterning in the epithelium and mesenchyme is well established, particularly through HOX gene expression, whether and how the ENS acquires region-specific transcriptional identities after settling into these niches remains an open question.

While the ENS transcriptional landscape remains less defined, the small intestine’s functional output exhibits clear regionalization along the anterior-posterior axis. Physiologically, motility patterns shift aborally: contraction frequencies step down from the duodenum to the ileum, and propulsive efficacy varies to accommodate the transition from rapid mixing to prolonged absorption. Structurally, recent work revealed that the ENS is organized into circumferential ‘stripes’ of varying neuronal density that differ along the length of the gut, potentially scaffolding these distinct motility gradients^4^. Furthermore, the neighboring epithelium is strictly zoned; a recent study demonstrated that the small intestinal epithelium is divided into five discrete metabolic domains, each governed by unique transcriptional programs^5^. Given these overt functional and epithelial boundaries within the small intestine, it is reasonable to hypothesize that the ENS possesses a parallel, regionally distinct transcriptional architecture.

The enteric nervous system (ENS) is a complex and autonomous network of neurons and glia embedded within the gastrointestinal tract. It regulates motility, secretion, absorption, local blood flow, and immune responses^6^. The ENS contains more neurons than the spinal cord and is organized into two major plexuses: the myenteric plexus, which regulates motility, and the submucosal plexus, which modulates secretion and vascular tone^7^. These neurons form intrinsic reflex circuits that sense mechanical and chemical cues within the gut lumen and coordinate appropriate responses. Derived from vagal neural crest (vNC) progenitors, ENS development involves extensive A-P migration, proliferation, and differentiation^8^. vNC cells originate from the caudal hindbrain, migrate through the somites, enter the foregut around E9 in mice, and then colonize the entire gastrointestinal tract^9^. This A-P migration imparts a temporal gradient to ENS assembly, but whether it also establishes enduring spatial specialization remains unresolved.

While early migratory and colonization events are well characterized, the later stages of ENS development remain far less understood, including the interplay between ENS progenitors and local tissue compartments, the transcriptional mechanisms of neuronal and glial diversification, and the establishment of neurotransmitter specification, axon trajectories, and circuit organization^10,11^. Classic studies have shown that Enteric Neural Crest Cells (ENCCs) require pathways such as Gdnf/Ret and Edn/Ednrb for colonization and survival^12,13^. However, it is unknown whether ENS cells subsequently adopt region-specific transcriptional programs that parallel the strong A-P identities of surrounding epithelial and mesenchymal tissues, or whether they instead maintain a largely uniform intrinsic program that is fine-tuned by local environmental cues and patterns of neuronal connectivity.

To dissect the relationship between gut regionalization and ENS development, we integrated multiplexed single-cell RNA sequencing (MULTI-seq) and functional perturbations. This combined approach allowed us to compare the spatial patterning of epithelium and mesenchyme with that of the ENS across developmental time, and to determine whether ENS cells acquire distinct regional identities or instead follow a predominantly temporal differentiation trajectory. By aligning ligand-receptor expression patterns with spatially resolved transcriptomics, we further investigated whether localized microenvironmental cues from the mesenchyme contribute to subtle transcriptional tuning in the ENS despite the absence of overt intrinsic regionalization. This framework enabled us to evaluate both the extent of A-P patterning within each compartment and the mechanisms through which extrinsic signals may refine ENS maturation within the context of a strongly regionalized gut.

We find that the epithelial and mesenchymal compartments display pronounced and persistent A-P patterning, including in expression of Hox genes and presence of region-specific cell types. In contrast, the ENS maintains a uniform vagal Hox identity and is primarily organized along a temporal maturation axis. Despite the absence of strong intrinsic regionalization, region-specific mesenchymal ligands, including PTPRZ1 ligands PTN and MDK, correlate with subtle ENS transcriptional shifts. Functional perturbations in human pluripotent stem cell-derived ENS cultures confirm that these microenvironmental cues regulate ENS progenitor proliferation, neuronal maturation, and neurotransmitter specification. Together, these findings support a model in which the ENS retains a stable core neuroglial program that is secondarily refined by localized, region-specific niche signals within the patterned gut.

## RESULTS

### The gut is strongly regionalized along the A-P axis, but the ENS is largely spatially uniform

We previously generated a spatiotemporal single-cell atlas of mouse small intestine (MSI) spanning E13.5–E18.5 where we segmented the tissue along the A-P axis at 2.5 mm spatial resolution (Fig. 1A)^14^. These samples were multiplexed from a single gut to minimize batch effects, with multiple experiments providing data across this developmental window. To better understand the detailed transcriptional events controlling ENS patterning we refined the resolution of this data set by integrating across multiple experiments using CONCORD^14^. The resulting unified latent space was well integrated across experimental data sets yet revealed the expected transcriptional differences across developmental time (Fig. 1B, Table 1).

**Figure 1.**
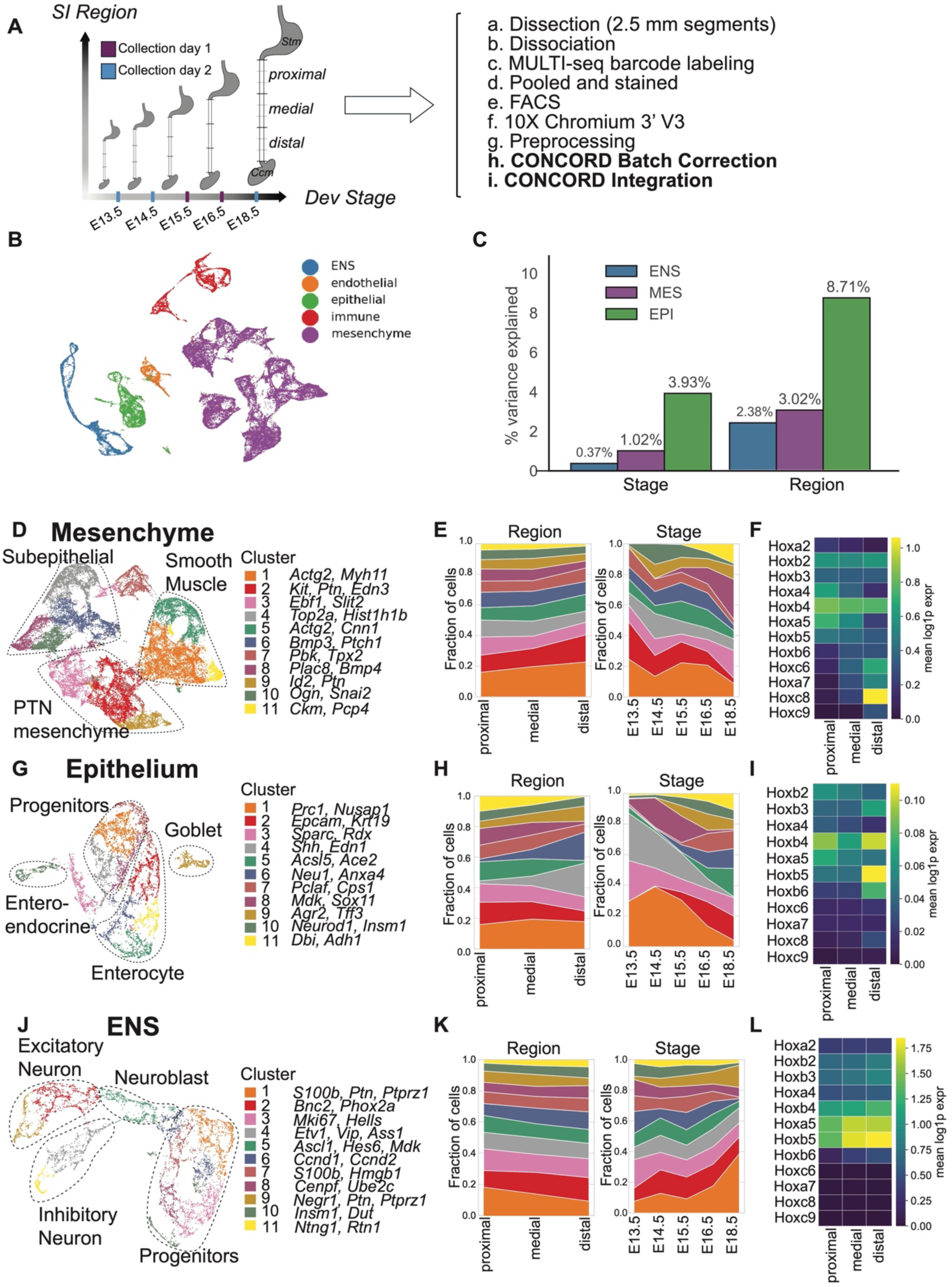
The developing gut is strongly regionalized along the A-P axis, but the ENS is largely spatially uniform. A: Schematic of the spatiotemporal single-cell atlas of the developing mouse small intestine B: UMAP of all MSI cells colored by broad compartment annotation (ENS, mesenchyme, epithelium, endothelium, immune). C: Bar plot showing the percentage of variance in gene expression explained by developmental stage and region for the ENS, mesenchyme (MES), and epithelium (EPI) compartments. D: UMAP of the MES subset colored by leiden clusters. E: Fraction of MES clusters across regions and developmental stages. F: Heatmap showing the mean expression of Hox genes along the anterior-posterior (AP) axis in the MES G: UMAP of the EPI subset colored by leiden clusters. H: Fraction of EPI clusters across regions and developmental stages. I: Heatmap showing the mean expression of Hox genes along the anterior-posterior (AP) axis in the EPI J: UMAP of the ENS subset colored by leiden clusters. K: Fraction of ENS clusters across regions and developmental stages. L: Heatmap showing the mean expression of Hox genes along the anterior-posterior (AP) axis in the ENS

To investigate the extent of patterning within each tissue compartment we applied variance partitioning using split-metadata PCA. We found that A-P position explained substantially more variance in epithelial (EPI) and mesenchymal (MES) than ENS, whereas developmental time contributed comparably to all (Fig. 1C, Table 2). Subsetting this atlas yielded a high-resolution MES compartment encompassing known fibroblast- and smooth-muscle derived subtypes (Fig. 1D,E, Table 3, Fig. S1A). Hox expression patterns revealed AP-graded signatures in MES (Fig. 1F). The EPI compartment also revealed AP-graded Hox expression (Fig. 1I) and additional cellular diversity including enterocytes, goblet cells, and enteroendocrine cells (Fig. 1G,H, Table 4, Fig. S1B). ENS cells formed clear progenitor, neuronal, and glial identities based on canonical markers (Sox10 for progenitors; Elavl4 for neurons; S100b for glia like progenitors, Fig. 1J,K; Fig. S1C; Table 5). In contrast to the epithelial and mesenchymal cells, Hox gene expression analysis showed that ENS retained a largely uniform vagal Hox profile with minimal regional variation (Fig. 1L). ENS Hox gene expression was stable over time [[[figure]]]. To investigate patterning and cellular composition across regions and stages, we quantified the proportion of each cluster across the A-P axis and developmental time. MES and EPI displayed clear A-P partitioning and sharper stratification by developmental age and region (Fig. 1E,H; Fig. S1D, E). In contrast, projecting A-P position and developmental stage onto the ENS UMAP revealed no substantial differences in regional cell composition (Fig. 1K; Fig. S1F). Quantification of ENS subtype proportions showed overall stability across A-P regions and stages, aside from modest increases in two PTN⁺/PTPRZ1⁺ clusters (clusters 1 and 9; Fig. 1K). MES and EPI, on the other hand, exhibited both changes in cell-type proportion and clusters that emerge at specific time points (Fig. 1E,H). Combined, these data indicate that ENS transcriptional program is shaped primarily by temporal maturation rather than A-P regionalization.

### Microenvironmental signals, including PTN/MDK, provide potentially spatially structured inputs to the ENS

Given that the ENS lacks a robust intrinsic transcriptional code for A-P patterning, we hypothesized that regional specialization is instead imposed by extrinsic cues from the highly patterned gut microenvironment. To systematically identify position-dependent signals capable of instructing ENS development, we modeled intercellular communication between the ENS and its surrounding niches using CellChat (Fig. 2A). Mesenchyme emerged as the dominant source of incoming ligand-receptor signals to ENS across all stages and regions, with ENS autocrine interactions contributing secondarily (Fig. 2B). Pathway-level contrasts revealed a restricted set of spatially or temporally regulated signaling families, including PTN, MDK, EDN, NOTCH, and NCAM (Fig. 2C,D). Among these, PTN and MDK ranked consistently high across A-P regions and developmental stages, showing coherent spatial and temporal trends (Fig. 2E-G, Table 6). To identify the transcriptional basis for these effects, we modeled ligand and receptor expression as a function of position and developmental stage (Fig. 2H, I, Tables 7-9). At E15.5, PTN ligand exhibited strong spatial gradients within MES, EPI, and ENS (Fig. 2J,K; Fig. S2A-D), while the receptor PTPRZ1 showed one of the most pronounced spatial slopes within ENS (Fig. 2P). Temporally, MDK decreased but PTN and PTPRZ1 expression increased within ENS, paralleling the rise of PTN-enriched subclusters (Fig. 2J, Fig. 1J; Fig. S2E-H). These analyses position PTN/MDK-PTPRZ1 signaling as a spatially graded, developmentally dynamic microenvironmental axis with potential activity on the ENS. PTPRZ1 (protein tyrosine phosphatase receptor type Z1) is a receptor-type tyrosine phosphatase that is catalytically inactivated upon ligand binding. When PTN or MDK bind to the extracellular domain of PTPRZ1, they induce receptor dimerization and allosterically inhibit its phosphatase activity., leading to increased tyrosine phosphorylation of downstream substrates and enhanced activity of signaling pathways normally suppressed by PTPRZ1, including Src, β-catenin, and MAPK cascades.

**Figure 2.**
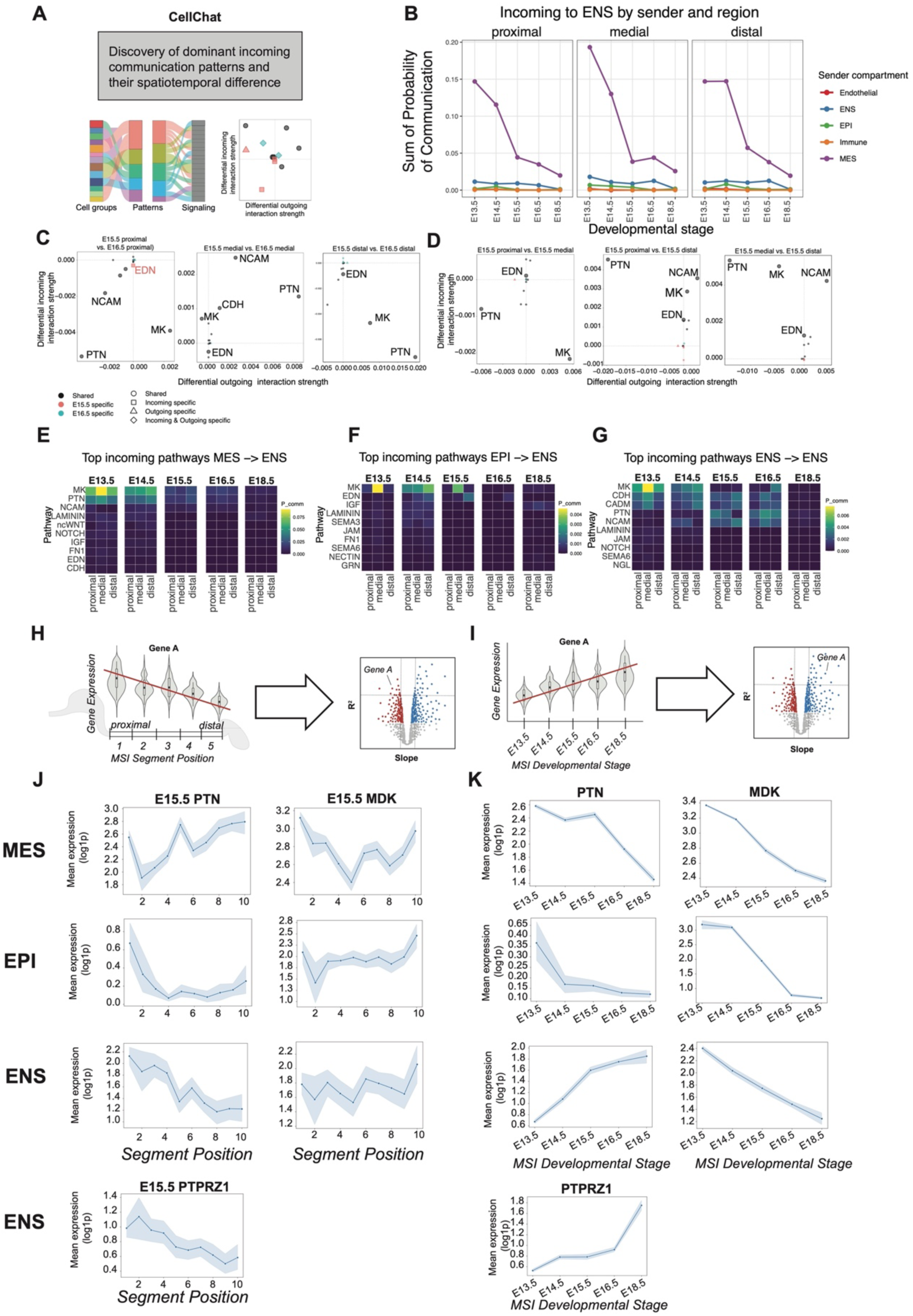
Microenvironmental signaling to ENS highlights PTN/MDK pathways with spatial and temporal structure. A: Conceptual schematic of the intercellular communication analysis using CellChat to model ligand–receptor (LR) communication across compartments. B: Line plot showing the net incoming signaling strength (Probability of Communication from CellChat) to the ENS from the sender compartments stratified by developmental stage and A-P region. MES is the dominant signaling source. C: Volcano plots summarizing pathway-level contrasts along the developmental axis (E15.5 proximal vs. E16.5 proximal), highlighting pathways like PTN, MDK, EDN, NCAM D: Volcano plots summarizing pathway-level contrasts along the A-P axis of MSI (distal vs. proximal) and temporal axis (E15.5 proximal vs. E15.5 distal), highlighting pathways like PTN, MDK, EDN, CDH, NCAM. E: Heatmap showing the strength of top incoming pathways from MES to the ENS across A-P regions and developmental stages. PTN and MDK have the highest probability of communication F: Heatmap showing the strength of top incoming pathways from EPI to the ENS across A-P regions and developmental stages. EDN and MDK signaling have the highest probability of communication. G: Heatmap showing the strength of top incoming pathways from ENS to the ENS across A-P regions and developmental stages. PTN and MDK signaling have the highest probability of communication H, I: Schematics of the spatial (H) and temporal (I) regression models used to link Ligand-Receptor transcript levels to position and age. J: Mean expression of Mdk and Ptn and their receptor Ptprz1 vs MSI segment position from the spatial regressions at E15.5 K: Mean expression of Mdk and Ptn and their receptor Ptprz1 vs MSI segment position from the temporal regressions

### PTN-expressing ENS subclusters represent distinct developmental states

Given the prominence of signatures for PTN signaling in these data, we examined two ENS subclusters highly enriched for PTN/MDK–PTPRZ1 components (Fig. S3): cluster 1 (Ptn⁺/Ptprz1⁺; Plp1⁺, S100b⁺) and cluster 9 (Ptn⁺/Ptprz1⁺; Elavl4⁺, Negr1⁺, Hoxb5⁺), both of which increased in abundance over developmental time (Fig. 1J, K). DEG analysis between cluster 1 and other progenitors (Fig. 3A) displayed a progenitor-like transcriptional signature, including translation at synapse level (Fig. 3B, C). Gene regulatory network analysis using SCENIChighlighted glial-biased or early progenitor regulons such as NFIX, FOS, and SOX10 (Fig. 3D). PTN- and PTPRZ1-centered GRNs consisted of TF hubs and targets associated with progenitor maintenance, cytoskeletal remodeling, and early lineage priming (Fig. 3E, F). Cluster 9 (Fig. 3G) demonstrated a mature neuronal-committed profile, with upregulation of axonogenesis, neurite extension, synaptic programs, and ion channels (Fig. 3H, I) in comparison to other neuronal clusters. Its regulon landscape differed markedly from cluster 1, showing increased activity of neuronal maturation TFs, including HOX family members, KLF7, and ZFHX2 (Fig. 3J-L). Together, these data reveal that PTN-enriched ENS populations encompass transcriptionally and regulatorily distinct developmental states, suggesting that PTN signaling may serve different roles in progenitor versus neuronal lineages.

**Figure 3.**
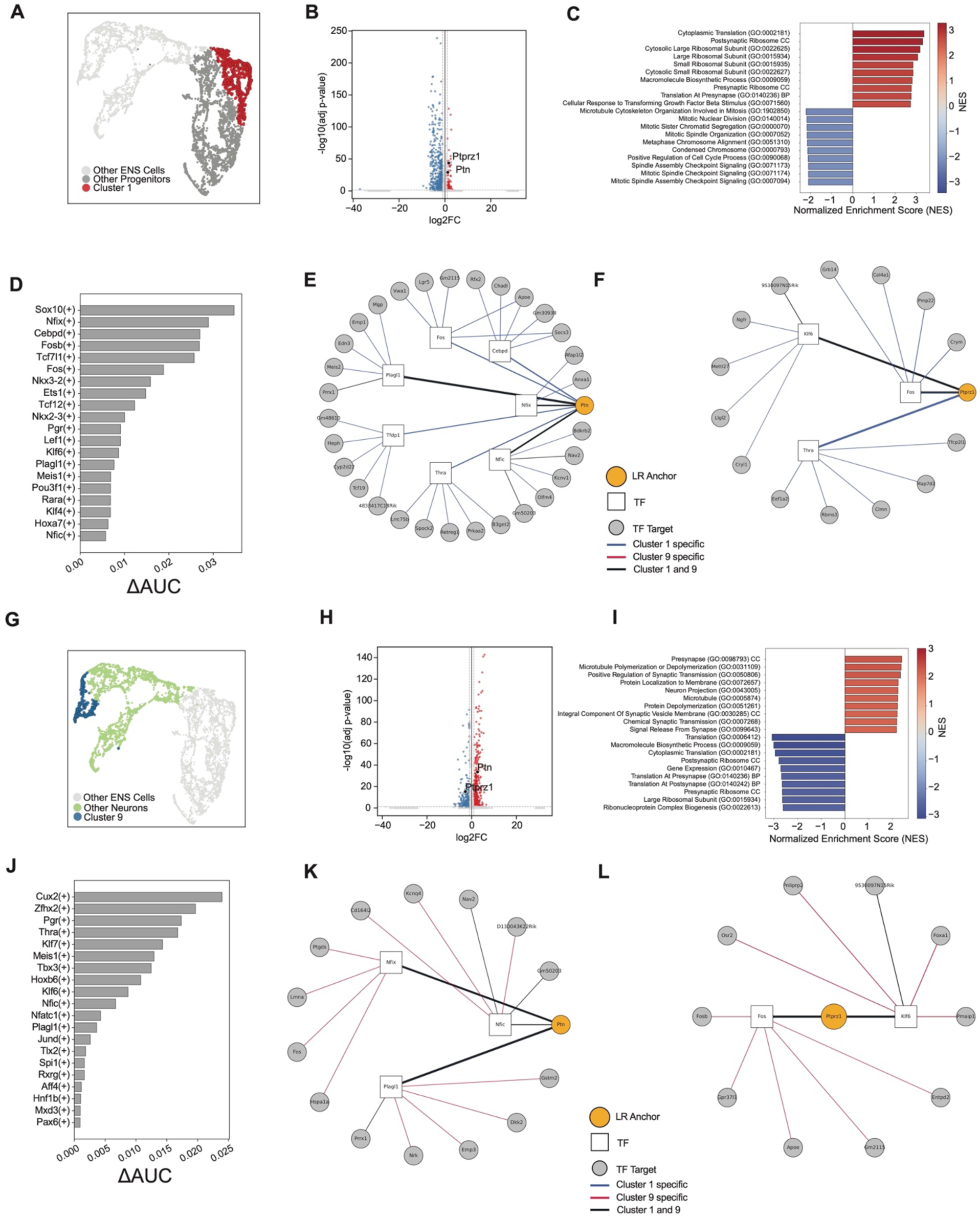
PTN-expressing ENS subclusters exhibit distinct developmental programs and gene regulatory networks. A: UMAP of the ENS subset highlighting progenitor cluster 1 with high Ptn and Ptprz1 expression. B: Volcano plot of DEGs between cluster 1 and other progenitors. C: GSEA for ranked genes from B, showing enrichment of top significant terms related to translation, transcription, and TGF-beta signaling D: The change in AUCell score of top TF regulons activities (pySCENIC AUCell) that distinguish cluster 1 amongst other progenitors. Showing high activity of regulons like Sox10, Nfix, and Fos. E, F: Gene regulatory networks (GRNs) centered on Ptn (F) and Ptprz1 (G) in cluster 1. G: UMAP of the ENS subset highlighting neuronal cluster 1 with high Ptn and Ptprz1 expression. H: Volcano plot of DEGs between 9 and other neuronal progenitors. I: GSEA for ranked genes from H, showing enrichment of top significant terms related to synaptogenesis, axonogenesis, and neurotransmission J: The change in AUCell score of top TF regulons activities (pySCENIC AUCell) that distinguish cluster 9 amongst other progenitors. Showing high activity of regulons like Sox10, Nfix, and Fos. K, L: Gene regulatory networks (GRNs) centered on Ptn (F) and Ptprz1 (G) in cluster 9.

### PTN/MDK-PTPRZ1 signaling modulates neurogenesis and neurotransmitter specification in hPSC-derived ENS

To test the functional relevance of these pathways, we perturbed the PTN/MDK-PTPRZ1 pathway in hPSC-derived ENS cultures. We used NAZ2329, which is a cell-permeable small molecule that binds to the active D1 catalytic domain of PTPRZ1 and stabilizes an open, inactive conformation of the WPD catalytic loop, to mimic the native signal.

Day 30 enteric ganglioids were treated with PTN, MDK, or the PTPRZ1 inhibitor NAZ2329 followed by bulk RNA-seq (Fig. 4A). PCA revealed condition-specific transcriptional shifts, with biological replicates clustering by treatment (Fig. 4B). PC1 loadings represent genes involved in mitosis and PC2 reflects extracellular matrix organization (see Tables 10-13). Across perturbations, ENS lineage programs were broadly altered (Fig. 4C-H). PTN, MDK, and NAZ2329 downregulated neuronal and synaptic maturation modules while upregulating progenitor and proliferation signatures. GSEA confirmed negative enrichment of axon development and synapse assembly, accompanied by positive enrichment of E2F, G2/M, and mitotic pathways (Fig. 4D).

**Figure 4.**
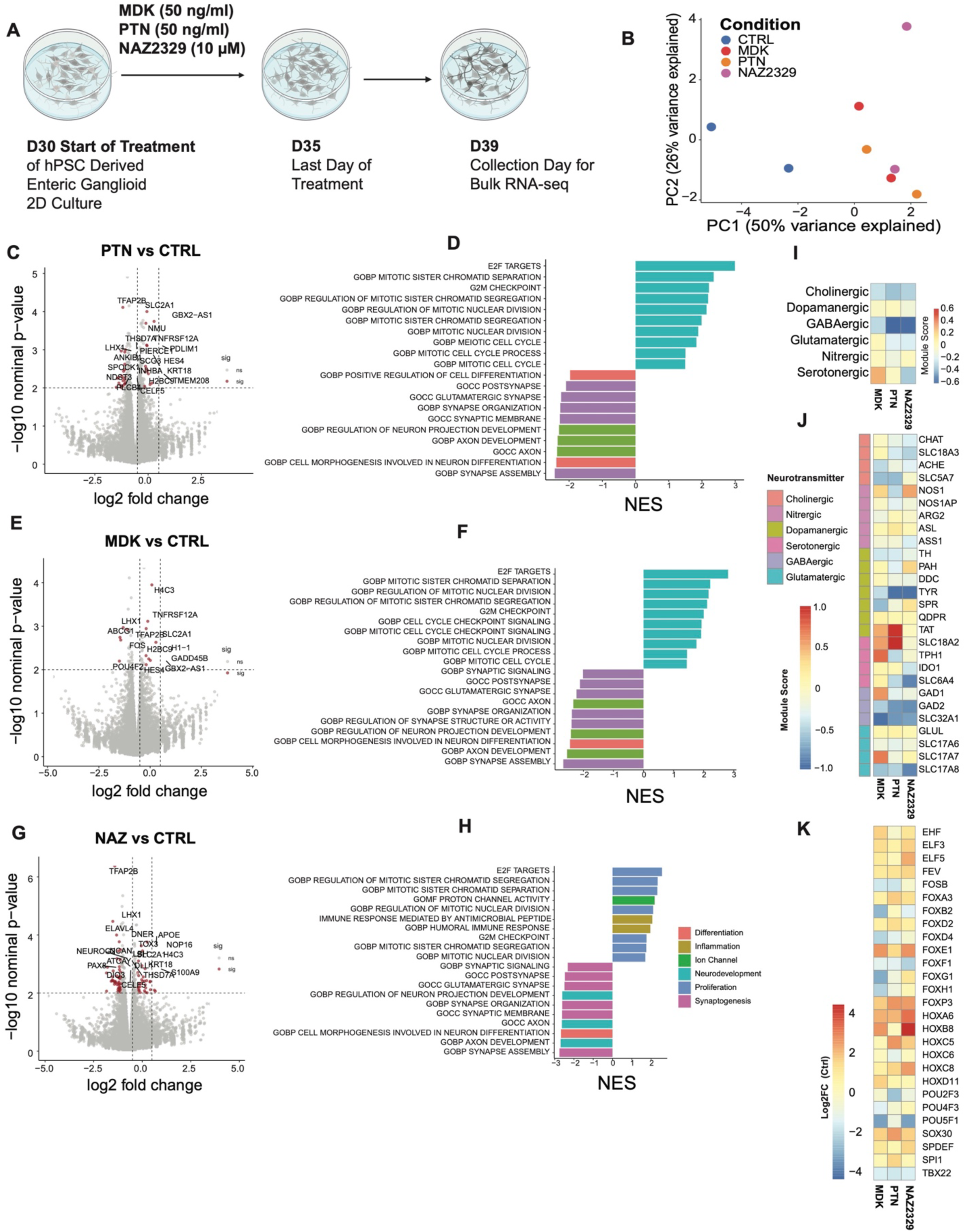
Ligand perturbations of PTN/PTPRZ1 and EDNRB axes reweight proliferative and synaptic maturation programs in human ENS cultures. A: Schematic of the bulk RNA-seq perturbation experiment on hPSC-derived ENS cultures. B: PCA of bulk RNA-seq samples, showing that samples cluster by treatment condition (CTRL, MDK, PTN, NAZ2329) C: Volcano plot showing the DEG of PTN relative to CTRL D: GSEA of ranked genes from C show positive enrichment of terms related to proliferation and negative enrichment of maturation terms E: Volcano plot showing the DEG of MDK relative to CTRL F: GSEA of ranked genes from C show positive enrichment of terms related to proliferation and negative enrichment of maturation terms G: Volcano plot showing the DEG of NAZ2329 relative to CTRL H: GSEA of ranked genes from C show positive enrichment of terms related to proliferation and negative enrichment of maturation terms I: Heatmap showing the log₂ fold-change in neurochemical module scores relative to CTRL. MDK increase neuronal serotonergic modules, while PTN and NAZ2329 downregulate GABAergic-associated modules. J: Heatmap showing the log₂ fold-change in neurochemical genes relative to CTRL. MDK and PTN increase neuronal serotonergic modules, while PTN, MDK, and NAZ2329 downregulate GABAergic-associated genes. K: Heatmap of log₂ fold-changes for selected TF genes. HOX and POU families had largest changes.

Treatment with PTPRZ1 inhibitors also changed the expression of neurotransmitter-identity modules (Fig. 3I, J). GABAergic identity was consistently suppressed, with PTN showing the strongest effect (reductions in GAD2, SLC32A1). PTN selectively increased serotonergic markers, including SLC18A2. Other neurotransmitter classes showed more modest or condition-specific responses. Inhibition of PTPRZ1 receptor via PTN, MDK, and NAZ2329 increased expression of HOX genes and decreased the expression of POU transcription factors. These perturbations also induced widespread remodeling of cytokine, neuropeptide, and receptor expression, indicating broader changes to signaling interactions (Supplementary Fig. 2A-J; Tables 14 and 15). These data indicate that PTPRZ signaling regulates proliferative, neurogenic, and neurotransmitter-specification transcriptional programs during human ENS differentiation, although, the observed transcriptional shifts reflect a combination of cell-intrinsic gene expression changes within individual cell types (e.g., progenitors, neurons, glia) and potential changes in the relative proportions of these cell types due to limitations of bulk RNA-seq.

Collectively, these results support a model in which the epithelium and mesenchyme establish strong A-P regional identities, whereas ENS cells maintain a largely uniform intrinsic program centered on temporal neuroglial maturation. Despite the absence of intrinsic ENS regionalization, local mesenchymal cues, particularly PTN/MDK family ligands, fine-tune ENS progenitor states, neuronal maturation trajectories, and neurotransmitter identity. This framework explains how ENS cells integrate into a highly regionalized organ while preserving a stable core identity and highlights localized niche signaling as a key mechanism for region-specific ENS refinement.

## DISCUSSION

In this study we find that gut epithelium and mesenchyme autonomously form strong A-P patterning, including a prominent region-specific HOX code. In contrast, this regional patterning is not mirrored within the ENS. Instead, ENS cells retain a largely uniform intrinsic transcriptional identity anchored in vagal neural crest origin and organized predominantly along a temporal maturation axis. This contrast indicates that ENS progenitors do not adopt local positional identities in the same manner as surrounding gut tissues, despite experiencing the same spatial environment.

While the ENS appears to lack intrinsic A-P regionalization, our analyses points to a variety of microenvironmental inputs to its development. Ligand-receptor profiling and spatial transcriptomics identify a small number of mesenchyme-derived signaling pathways, including PTN/MDK, EDN, and NOTCH, as spatially enriched and developmentally dynamic inputs to ENS cells. Among these, PTN/MDK–PTPRZ1 signaling emerges as a prominent regulator with combined spatial gradients and temporal changes that parallel ENS maturation. These signals correlate with subtle transcriptional tuning in ENS subpopulations, suggesting that local microenvironmental cues modulate ENS development even in the absence of strong intrinsic regionalization.

Functional perturbation of PTPRZ1 activity in human pluripotent stem cell–derived ENS cultures support this model. Perturbation of PTPRZ1 signaling alters ENS lineage trajectories by enhancing proliferative programs, suppressing synaptic and neuronal maturation modules, and shifting neurotransmitter specification, for example, reducing GABAergic identity while elevating serotonergic markers.

While the specific role for PTPRZ1 signaling is not addressed in this study, these findings point to a potential effect on PTPRZ1 signaling. Specifically, PTPRZ1 signaling in human ES-cell derived enteric neural progenitors balances progenitor expansion with neuronal differentiation, i.e., when PTPRZ1 is inhibited, the resulting enhancement of kinase signaling promotes a more proliferative, less differentiated state while simultaneously biasing neurotransmitter fate decisions toward specific identities such as serotonergic neurons.

These coordinated changes across transcriptional, regulatory, and signaling layers highlight how microenvironmental cues fine-tune ENS development. Importantly, these effects are observed across both mouse and human systems, underscoring the evolutionary conservation of these niche interactions.

Together, our findings support a model in which ENS cells establish a uniform neuroglial “core program” early in development that can be sculpted by localized, regionally patterned signals from the mesenchyme. This model reconciles two longstanding observations that the gut exhibit high regional specialization while the ENS is largely transcriptionally uniform^15,16^. While ENS cells do not encode overt A-P positional identities intrinsically, they appear to integrate spatial information through microenvironmental cues that fine-tune their maturation, connectivity, and neurotransmitter specification as they differentiate within distinct gut niches.

Our findings lead to several intriguing questions that warrant future investigations. One is whether ENS regionalization is further refined or emerges more strongly at later developmental or postnatal stages, potentially through activity-dependent maturation or epigenetic reprogramming. Another is how additional signaling axes beyond those identified here, such as PTN/MDK, EDN, NOTCH, and WNT, interact with or build upon early niche-derived cues to shape ENS lineage transitions and circuit formation over time. Finally, while our study identifies mesenchyme-derived ligands as key modulators of ENS maturation, the relative contributions of other microenvironmental features, including secreted factors, extracellular matrix components, metabolic gradients, and short-range cell–cell interactions to ENS development and function remain incompletely understood. How disruptions in these microenvironmental cues contribute to enteric neuropathies is an important question for future investigation.

By providing a multi-lineage view of gut development, our study defines how intrinsic programs interface with regionally patterned niches to produce a mature, spatially integrated ENS. This framework not only refines our understanding of ENS development but also has implications for regenerative strategies, disease modeling, and cell-based therapies, which must account for the role of microenvironmental context in shaping ENS function.

## METHODS

### Mouse embryonic small intestine scRNA-seq dataset

The single-cell RNA sequencing (scRNA-seq) dataset of embryonic mouse small intestine (MSI) analyzed in this study was generated previously and described elsewhere^17^. In brief, wild-type mouse embryos were collected at embryonic days (E) 13.5, 14.5, 15.5, 16.5 and 18.5. For each embryo, the small intestine was dissected from stomach to cecum in ice-cold PBS under a stereomicroscope. The MSI was then subdivided into contiguous 2.5 mm segments along the anterior-posterior (AP) axis. Each segment was assigned a segment position index (SegPos), and the total number of segments per embryo increased with developmental stage to approximate equal physical lengths. For some analyses, segments were subsequently grouped into proximal, medial, and distal regions by binning SegPos into thirds along the A-P axis. Embryo ID, SegPos, and derived region labels were retained as metadata throughout the analysis.

For each embryo, pooled segments were enzymatically and mechanically dissociated into single-cell suspensions as previously described^18^. Briefly, tissue was incubated in a digestion buffer containing collagenase/dispase and DNase I at 37 °C with gentle agitation, followed by trituration and filtration through a 40 µm strainer. Live cells were enriched either by flow cytometry or density-based methods. Single-cell suspensions from multiple segments and embryos were then multiplexed using MULTI-seq lipid-modified DNA oligonucleotide barcodes following the published protocol^19^. After barcode labeling, cells from multiple samples were pooled, optionally re-sorted to enrich for viable singlets, and loaded onto the 10x Genomics Chromium platform for single-cell 3′ RNA-seq library preparation according to the manufacturer’s instructions. Libraries were sequenced on an Illumina platform to a depth sufficient to achieve approximately tens of thousands of reads per cell.

Raw base call files were processed using Cell Ranger (10x Genomics) against the appropriate mouse reference genome (mm10) to generate gene-by-cell count matrices. MULTI-seq barcodes were demultiplexed as described previously^19^ to assign each cell to a single sample or to identify barcode collisions and likely doublets. Our analyses used a curated AnnData object generated from this dataset, in which initial quality control and demultiplexing had been performed. Low-quality cells were excluded based on minimum UMI counts, gene complexity, and mitochondrial read fraction; thresholds were tuned per batch and matched those used in the original study. Doublets were identified with Scrublet^20^ and by MULTI-seq barcode conflicts and removed. Genes expressed in very few cells were excluded prior to normalization.

### Normalization, batch integration, and clustering

All single-cell analyses were performed in Python using scanpy and anndata unless otherwise stated^21^. For each cell, raw counts were library-size normalized (for example, scaling total counts to 10,000 per cell) and log-transformed using the natural logarithm of counts plus one (log1p) to generate a log-normalized expression matrix. Highly variable genes (HVGs) were identified within each batch using scanpy’s HVG functions, and batch-specific HVGs were merged after excluding mitochondrial and ribosomal genes from consideration. To integrate data across developmental stages and embryos, we used CONCORD to learn a shared low-dimensional latent space^14^. Principal component analysis (PCA) was performed on the scaled HVG expression matrix, and k-nearest neighbor (kNN) graphs were constructed for each batch. CONCORD was then applied with default domain alignment hyperparameters to jointly align these graphs across batches and generate a unified latent representation for all cells. This latent embedding was stored in the AnnData object and used to compute a global kNN graph and a UMAP embedding^22,23^. All UMAP visualizations shown in Figures 1 and 3 were derived from this CONCORD-based neighbor graph and stored as .obsm[“Concord_UMAP”]. Cell clustering was performed using the Leiden community detection algorithm on the integrated kNN graph^24^. We explored a range of resolutions and selected those that yielded stable and biologically interpretable clusters. A global clustering solution was used to identify major compartments (enteric nervous system [ENS], mesenchyme [MES], epithelium [EPI], endothelium, immune), and independent clustering was then performed on compartment-specific subsets for higher resolution analyses. The resolution used for each subset is reported in the figure panels.

### Cell-type annotation and compartment subsetting

Broad cell types were annotated on the integrated atlas by inspecting canonical marker genes and, where appropriate, referencing annotations from the original study^17^. ENS cells were identified by expression of Sox10, Ednrb, Ret, Phox2a, Phox2b and related markers. Mesenchymal cells were defined by Pdgfra, Col1a1, Vim and other mesenchymal markers; epithelial cells by Epcam and keratins; endothelial cells by Pecam1 and Kdr; and immune cells by Ptprc and lineage-specific genes (see Table 1-4). For downstream analyses, we created separate AnnData subsets for ENS, MES and EPI compartments by filtering on these broad labels. Each subset retained CONCORD latent and UMAP coordinates, Leiden cluster identities, SegPos, region assignment, Age, embryo ID and QC covariates.

### Composition across developmental stage and A-P region

To quantify how ENS, MES, and EPI subtypes are distributed across developmental stages and A-P regions, we computed cluster proportions at the embryo level. For each embryo and condition (for example, a particular stage or region), we counted the number of cells belonging to each Leiden cluster within a given compartment and divided by the total number of cells from that compartment in the same embryo and condition. These embryo-level proportions were then aggregated by taking the mean across embryos for each condition and visualized as stacked bar or area plots (Figure 1I-L). To derive uncertainty estimates, we performed nonparametric bootstrap resampling of embryos within each condition and recomputed proportions to obtain 95% confidence intervals. Where formal statistical testing of compositional changes was required, generalized linear models with binomial or Dirichlet-multinomial likelihoods were fitted to embryo-level counts, with condition as the main predictor^25^.

### Variance partitioning

To compare the relative contributions of developmental stage and A-P region to transcriptional variance within ENS and MES, we implemented a split-metadata PCA framework. For each compartment, log-normalized HVG expression was scaled, and PCA was performed. For each principal component (PC), we fitted a linear model regressing PC score on Age, region, cell-cycle covariates and batch. From these models, we extracted partial eta-squared values for each factor, representing the fraction of variance in that PC explained uniquely by that factor conditional on the others. We then aggregated partial eta-squared values across PCs by weighting them by the proportion of total variance explained by each PC. This yielded compartment-specific estimates of the variance attributable to stage and region, which were visualized with bootstrapped confidence intervals in Figure 1C (See summary Table 2).

### Ligand-receptor communication analysis with CellChat

Intercellular communication between compartments was inferred using CellChat (R package), focusing on signaling pathways targeting ENS^26^. Raw counts and metadata from the integrated atlas were exported to R, and CellChat objects were constructed after filtering for relevant sender and receiver groups. ENS, MES, and EPI were included as both potential senders and ENS receivers, along with endothelial and immune compartments. For each Age and region combination, we derived a communication network using the CellChatDB.mouse database of ligand-receptor pairs. Overexpressed genes and interactions were identified using CellChat’s standard pipeline, communication probabilities were computed for each LR pair, and pathway-level communication probabilities were aggregated by summing over LR pairs belonging to the same signaling family. Net incoming signaling strength to ENS was defined as the sum of pathway-level communication probabilities for all interactions with ENS as the receiver, broken down by sender compartment, stage, and region. These values were visualized in Figure 2B to highlight the dominant contribution of EPI- and MES-derived signals, followed by ENS autocrine interactions. To identify pathways with spatial or temporal signatures, we compared pathway-level “information flow” metrics along the A-P axis and across development. For spatial contrasts (for example, distal vs proximal), we computed differences in information flow and assessed significance by permutation testing of condition labels (typically several thousand permutations per comparison), followed by Benjamini-Hochberg correction for multiple pathways. An analogous strategy was used for temporal contrasts (for example, E13.5 vs E14.5). Volcano plots in Figure 2C-D summarize standardized differences in information flow versus –log10(FDR), highlighting pathways such as PTN, MDK, EDN, NOTCH. For selected top pathways, including PTN, we also plotted pathway strength across regions and stages with bootstrapped confidence intervals.

### Spatial and temporal regression of ligand and receptor expression

To link pathway-level CellChat results to underlying transcript dynamics, we modeled expression of individual ligands and receptors as functions of A-P position and developmental stage. For each gene of interest, we considered log-normalized expression in the relevant compartment (for example, EPI, MES, ENS ligands, ENS receptors) and fitted linear models at the single-cell level. Spatial models related expression to numeric segment position (SegPos) within a fixed developmental stage, including covariates for batch, cell-cycle score, total UMI counts and mitochondrial fraction. Temporal models related expression to Age across stages, with region included as a covariate alongside the same technical and biological covariates. From each model, we obtained regression slopes, coefficients of determination (R²), and p-values for the variable of interest (SegPos or Age). P-values were adjusted across genes using Benjamini-Hochberg. Genes were visualized on volcano plots with slope on the x-axis and R² on the y-axis, and those with significant trends and high explanatory power were plotted. For top candidates such as Ptn and Ptprz1, we also plotted mean expression across ordered segments or developmental stages with bootstrapped 95% confidence intervals to illustrate spatial gradients and temporal trajectories in ligand and receptor expression.

### ENS subcluster definition and differential expression

Within the ENS subset, we identified subclusters with strong PTN/PTPRZ1 expression by overlaying Ptn and Ptprz1 expression on the ENS UMAP and inspecting Leiden clusters. Two clusters were selected for detailed analysis: a Ptn+/Ptprz1+/Plp1+ progenitor-like cluster (cluster 1) and a Ptn+/Ptprz1+/Elavl4+ neuronal-committed cluster (cluster 9). Cluster 1 was compared to other ENS progenitor-like clusters, whereas cluster 9 was compared to other ENS neurons. These focal and reference groups are shown in Figure 3A and 3G. Differential expression between each focal cluster and its corresponding reference group was performed using scanpy’s rank_genes_groups function^27^. Unless otherwise indicated, we used the Wilcoxon rank-sum test on log1p-normalized expression values and restricted analysis to genes expressed in at least a minimal fraction of cells in either group. For each gene, we obtained log₂ fold-changes and adjusted p-values (Benjamini–Hochberg). Genes were visualized on volcano plots (Figure 3B, 3H) with log₂ fold-change versus –log10(FDR), and significantly differentially expressed genes above an effect-size threshold were labeled. These DE signatures were then used as ranked lists for downstream gene set enrichment analysis.

### Gene set enrichment analysis for ENS subclusters

To interpret differential expression profiles for clusters 1 and 9, we performed gene set enrichment analysis (GSEA) using gseapy^28^. Genes were ranked by a signed statistic derived from the DE results (for example, log₂ fold-change weighted by significance), and this ranked list was queried against MSigDB Hallmark and Gene Ontology (biological process and cellular component) gene sets. Gene set definitions were obtained via msigdbr and harmonized with mouse gene symbols by case conversion and orthology mapping where necessary. Normalized enrichment scores (NES) and FDR-adjusted q-values were computed using permutation-based tests. Top 10 enriched pathways with FDR < 0.05 were summarized in bar plots (Figure 3C, 3I) and highlighted key differences between progenitor-like and neuronal-committed PTN+/Ptprz1+ states, including enrichment of mitotic, transcription, and translation programs at synapse in cluster 1 and neurite extension, synaptic pathways, and neurotransmission in cluster 9.

### Regulatory network inference and PTN/PTPRZ1-centered GRNs

Transcription factor (TF) regulons and gene regulatory networks were inferred using the pySCENIC workflow applied to the ENS subset^29^. Co-expression modules were first identified using GRNBoost2, using a curated list of mouse TFs as candidate regulators. These co-expression relationships were then filtered based on motif enrichment and cis-regulatory evidence using RcisTarget and species-specific motif databases, resulting in regulons defined as TFs and their direct target gene sets. Regulon activity for each cell was quantified with AUCell, which computes the area under the recovery curve of regulon genes in each cell’s ranked expression profile, yielding regulon activity scores (AUCs). To identify regulons distinguishing clusters 1 and 9 from their respective reference groups, regulon AUCs were compared using linear models or rank-based tests with cluster identity as the primary predictor and stage and region as covariates. Regulons with significant differences after FDR correction were selected and visualized as barplot (Figure 3D, 3J), showing patterns of regulon activity across clusters and developmental states. To construct PTN- and PTPRZ1-centered gene regulatory subnetworks, we extracted TF–target edges from the pySCENIC-inferred networks that involved PTN or PTPRZ1 either as TF targets or as TFs with strong regulon activity in PTN+/Ptprz1+ clusters. For each edge, we derived a composite weight by averaging normalized co-expression importance and normalized motif enrichment scores. We then selected TFs with high out-degree or cumulative edge weights within cluster 1 or cluster 9 and visualized these subnetworks (Figure 3E–F, 3K–L). The edge thickness represented composite weights, highlighting distinct TF hubs and downstream programs associated with PTN/PTPRZ1 in progenitor-like versus neuronal-committed ENS cells.

### hPSC maintenance, ENS differentiation, and perturbation design

Human pluripotent stem cells (hPSCs) were maintained in a defined medium (for example, mTeSR1 or Essential 8) and passaged using non-enzymatic dissociation reagents according to standard protocols. Cells were routinely tested for mycoplasma contamination. Differentiation toward enteric neural crest and ENS-like ganglioid cultures followed a previously established protocol^30–32^. In brief, hPSCs were first induced toward neural crest using combinations of dual-SMAD inhibition and Wnt activation, then patterned toward a vagal ENS fate using GDNF over the course of approximately two to three weeks. Cultures were then maintained in conditions that support ENS-like neuronal and glial maturation, producing dense networks of ENS-like ganglia by around Day 30 (D30) of differentiation. To interrogate PTN/PTPRZ1 and EDNRB signaling axes, Day-30 ENS ganglioid cultures were treated with recombinant ligands and small-molecule inhibitors. Cultures were exposed to midkine (MDK; 50 ng/mL), pleiotrophin (PTN; 50 ng/mL) or the small-molecule PTPRZ1 inhibitor NAZ2329 (NAZ; 10 µM). In parallel, cells were treated with an EDNRB agonist (EDN3) and an EDNRB antagonist (BQ), using concentrations previously shown to modulate ENS lineage behavior. Treatments were administered in neuronal induction/basal medium from D30 to D35, with media changes at appropriate intervals. On D35, treatments were discontinued, and cultures were maintained in basal medium until Day 39 (D39) to allow a brief chase period. All conditions, including vehicle-treated controls (CTRL), were performed with three independent biological replicates, defined as separate differentiation batches or independently seeded wells.

### RNA extraction, bulk RNA-seq library preparation, and sequencing

On D39, cultures were washed with PBS and lysed directly in RNA extraction buffer. Total RNA was isolated using a column-based kit according to the manufacturer’s instructions, including on-column DNase treatment to remove genomic DNA. RNA quality and integrity were assessed by TapeStation or Bioanalyzer, and concentration was measured by Qubit. Only samples with sufficient yield and acceptable RNA integrity were used for library preparation. Bulk RNA-seq libraries were prepared by Plasmidsaurus using Oxford Nanopore Technology with custom analysis and annotation. Raw sequencing reads were processed using a standard alignment and counting pipeline. Reads were adapter- and quality-trimmed, aligned to the human reference genome (GRCh38) using STAR^33^, and assigned to genes using featureCounts with GENCODE gene annotations^34^. This yielded a sample-by-gene count matrix that was imported into R for differential expression analysis with DESeq2.

### DESeq2 analysis, PCA, and sample-level QC

Differential expression analysis of bulk RNA-seq data was carried out in R using DESeq2^35^, with additional visualization performed using tidyverse and related packages. Raw counts and sample metadata (condition, replicate) were used to construct a DESeqDataSet, and size factors and dispersions were estimated using DESeq2’s standard workflow. A design formula with condition as the main factor was used to model expression differences among CTRL, MDK, PTN, and NAZ2329. For visualization, we applied the variance-stabilizing transformation (VST) to generate homoscedastic expression values. Principal component analysis (PCA) was performed on the VST-transformed matrix using prcomp. In Figure 4B, samples are plotted in the PC1–PC2 plane, colored by condition, showing separation of treatment groups and clustering of replicates. To aid interpretation and assess replicate consistency, condition centroids in PC space were computed and optionally overlaid. Outlier samples, defined by large distances from their condition centroid or inconsistent clustering based on correlation metrics, were identified in a separate QC script. Where outliers were detected, they were excluded from downstream DESeq2 contrasts used in figure panels; this filtering step is documented in Supplementary Methods. Differential expression contrasts were computed between each treatment and control (MDK vs CTRL, PTN vs CTRL, NAZ2329 vs CTRL, EDN vs CTRL, BQ vs CTRL) using DESeq2’s Wald test. For each gene in each contrast, DESeq2 reported log₂ fold-change, standard error, Wald statistic and adjusted p-value after Benjamini–Hochberg correction. Genes with FDR < 0.05 and |log₂ fold-change| above a nominal threshold were considered significantly differentially expressed.

### Module-based summaries and GSEA for bulk RNA-seq

To connect bulk perturbation responses to ENS developmental and neurotransmitter programs identified in the mouse scRNA-seq data, we defined curated gene modules representing progenitor, neuroblast, pan-neuronal, neuronal subtype and glial states, as well as neurotransmitter classes (for example, cholinergic, nitrergic, serotonergic, GABAergic) and signaling-related genes (PTN/PTPRZ1/EDNRB and downstream TFs). Module membership was based on literature and on cluster-specific markers from the mouse ENS atlas and is listed in Supplementary Tables 14 and 15. For each module and each DESeq2 contrast, we summarized treatment effects by computing the mean log₂ fold-change of genes within the module relative to CTRL. These module-level log₂ fold-changes were visualized as heatmaps (Figure 4C, 4E), with rows corresponding to modules or selected marker genes and columns corresponding to conditions. Where the goal was to compare absolute magnitudes across conditions, colors represent raw log₂ fold-changes. To obtain pathway-level insights, we performed gene set enrichment analysis on the bulk RNA-seq DESeq2 results using GSEA^36^ and fgsea^37^ and msigdbr^38^. For each contrast, genes were ranked by a signed statistic (for example, log₂ fold-change multiplied by –log10(FDR), or the DESeq2 Wald statistic), and this ranking was compared against MSigDB Hallmark and GO gene sets for Homo sapiens. Normalized enrichment scores and FDR-adjusted q-values were computed by fgsea. We focused on gene sets related to cell cycle and proliferation (for example, E2F targets, G2/M checkpoint, mitotic nuclear division), neurodevelopment and axon guidance, and synaptic structure and function. Figure 4D, 4F, 4H presents barplots of normalized enrichment scores for a curated subset of these pathways across treatments. This analysis revealed distinct and sometimes opposing effects of PTN/MDK/PTPRZ1 versus EDNRB signaling perturbations on proliferative and synaptic maturation programs in hPSC-derived ENS cultures. Category-specific heatmaps of log₂ fold-changes for neurotransmitter synthesis genes, neuropeptides, cytokines and receptors were also constructed to illustrate perturbation-induced changes in neurochemical identity and cell–cell communication components (Supplementary Figure 2), using curated gene lists and the same DESeq2-derived log₂ fold-changes.

### Statistics and software

Unless otherwise indicated, all statistical tests were two-sided, and multiple comparisons were controlled using the Benjamini–Hochberg false discovery rate procedure^39^. For single-cell analyses, effect sizes are reported as partial eta-squared for variance partitioning, regression slopes and R² values for spatial and temporal models, log₂ fold-changes for differential expression, and normalized enrichment scores for GSEA. For bulk RNA-seq, effect sizes are reported as DESeq2 log₂ fold-changes and normalized enrichment scores. Confidence intervals, where shown, represent 95% intervals derived from bootstrap resampling or model-based estimates. Single-cell analyses were implemented primarily in Python using scanpy^27^, anndata, numpy, pandas, scikit-learn, gseapy^28^, pySCENIC^29^ and custom scripts. CellChat^40^, DESeq2^41^, msigdbr^42^, GSEA^36^, tidyverse and heatmap packages (such as pheatmap or ComplexHeatmap) were used in R for ligand–receptor communication, bulk RNA-seq differential expression and pathway analyses.

## Acknowledgements

We thank all members of the Gartner and Fattahi laboratories, specifically Dr. Homa Majd, for their thoughtful feedback and discussions throughout the course of this study. We are grateful to Plasmidsaurus for Bulk RNA-seq library preparation and sequencing using Oxford Nanopore Technology, with custom analysis and annotation. This work was supported by the NIH Director’s New Innovator Award (DP2HD101401) to F.F., the California Institute for Regenerative Medicine Discovery Awards (DISC0-14521 and DISC2-15119) to F.F., NIH R01DK126376 (to Z.J.G.), and additional support from the Arc Institute Innovation Ignite Award to F.F. This work was also supported by the University of California, San Francisco (UCSF) Center for Cellular Construction, a National Science Foundation Science and Technology Center (DBI-1548297 to Z.J.G.). Z.J.G. is an investigator of the Chan Zuckerberg Biohub San Francisco.

---

**Figure S1.**
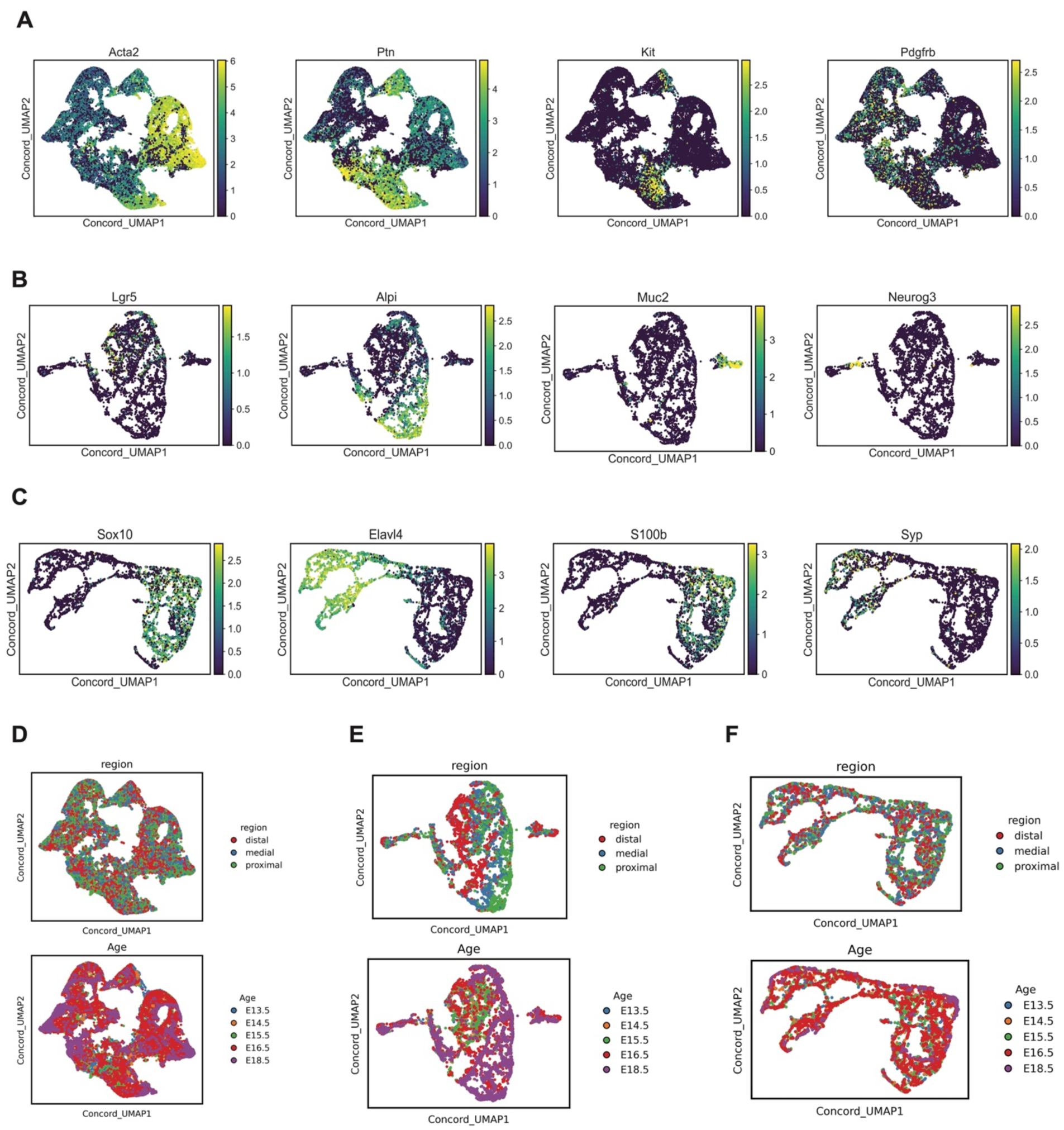
A: UMAP of the expression canonical markers in MES B: UMAP of the expression canonical markers in EPI C: UMAP of the expression canonical markers in ENS A: UMAP of the MES subset, with cells colored by MSI region and developmental stage. B: UMAP of the EPI subset, with cells colored by MSI region and developmental stage. C: UMAP of the ENS subset, with cells colored by MSI region and developmental stage.

**Figure S2:**
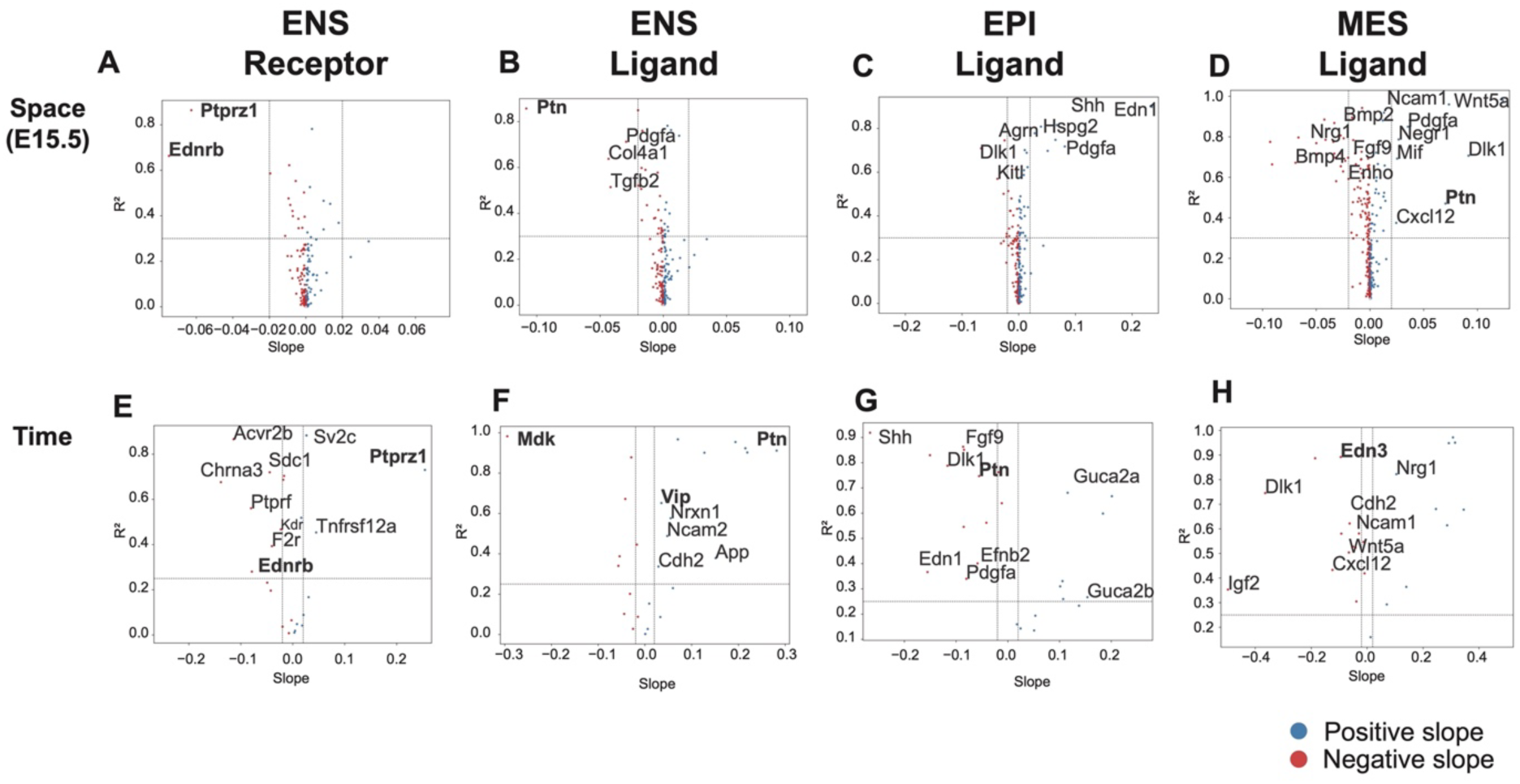
Volcano plots of the spatial (A-D) and temporal (E-H) regression models for ENS Receptors (A, E), ENS ligands (B, F), EPI ligands (C, G), MES ligands (D, H)

**Figure S3:**
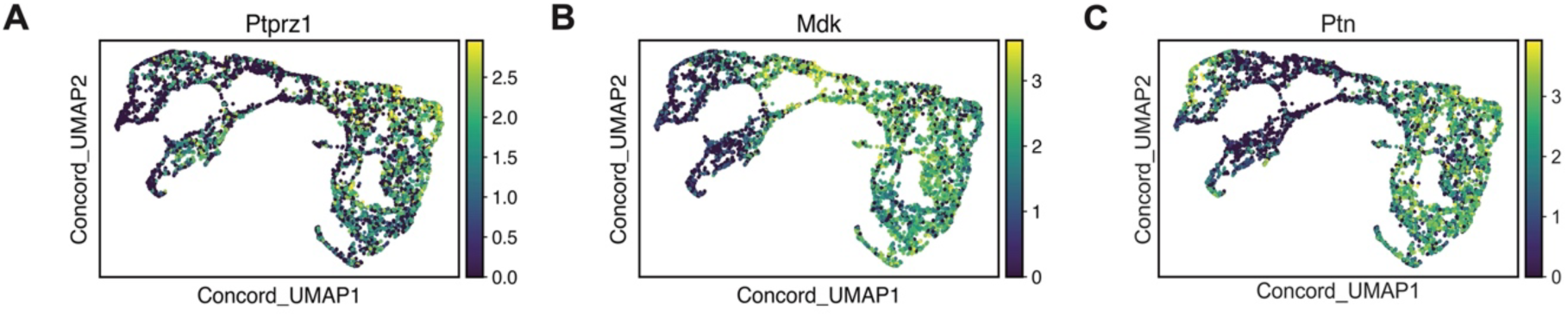
Expression of PTPRZ1 signaling components on Concord UMAP. A, B, C: UMAP highlighting the ENS cells with expression of (log₂ of counts) Ptprz1, Mdk, and Ptn.

**Figure S4:**
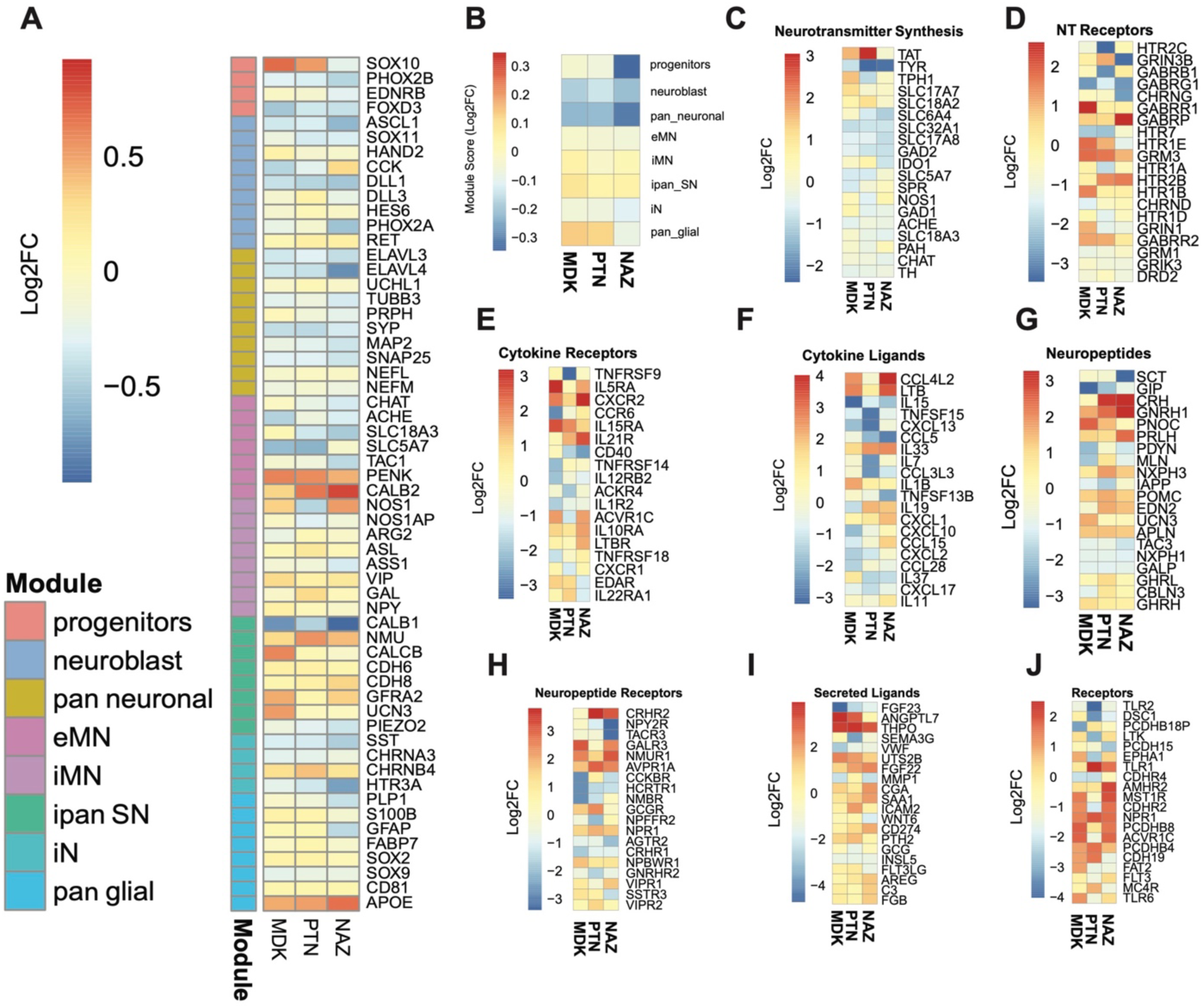
Transcriptomic consequences of PTPRZ1 inhibition via MDK, PTN, and NAZ2329. A: Heatmap showing the log₂ fold-changes in ENS state genes relative to CTRL. MDK, PTN, and NAZ2329 increase APOE, PENK, CALB2. MDK and PTN also increase SOX10 levels. B: Heatmap showing modest log₂ fold-changes in module scores relative to CTRL. MDK and PTN increase glial module, while NAZ2329 downregulate neuronal-associated modules. C: Heatmap showing modest log₂ fold-changes in neurotransmitter synthesis genes relative to CTRL. D: Heatmap showing modest log₂ fold-changes in neurotransmitter receptor genes relative to CTRL. E: Heatmap showing modest log₂ fold-changes in cytokine receptor genes relative to CTRL. F: Heatmap showing modest log₂ fold-changes in cytokine ligands relative to CTRL. G: Heatmap showing modest log₂ fold-changes in neuropeptide genes relative to CTRL. H: Heatmap showing modest log₂ fold-changes in neuropeptide receptor genes relative to CTRL. I: Heatmap showing modest log₂ fold-changes in secreted ligands relative to CTRL. J: Heatmap showing modest log₂ fold-changes in receptor genes relative to CTRL.

## REFERENCES

1. Wells, J. M. & Spence, J. R. How to make an intestine. Dev. Camb. Engl. 141, 752–760 (2014).

2. Zorn, A. M. & Wells, J. M. Vertebrate endoderm development and organ formation. Annu. Rev. Cell Dev. Biol. 25, 221–251 (2009).

3. Nagy, N. & Goldstein, A. M. Enteric nervous system development: A crest cell’s journey from neural tube to colon. Semin. Cell Dev. Biol. 66, 94–106 (2017).

4. Dershowitz, L. B. & Kaltschmidt, J. A. Enteric Nervous System Striped Patterning and Disease: Unexplored Pathophysiology. Cell. Mol. Gastroenterol. Hepatol. 18, 101332 (2024).

5. Zwick, R. K. et al. Epithelial zonation along the mouse and human small intestine defines five discrete metabolic domains. Nat. Cell Biol. 26, 250–262 (2024).

6. Furness, J. B. The enteric nervous system and neurogastroenterology. Nat. Rev. Gastroenterol. Hepatol. 9, 286–294 (2012).

7. Furness, J. B. The enteric nervous system: normal functions and enteric neuropathies. Neurogastroenterol. Motil. 20 Suppl 1, 32–38 (2008).

8. Le Douarin, N. M. & Teillet, M. A. The migration of neural crest cells to the wall of the digestive tract in avian embryo. J. Embryol. Exp. Morphol. 30, 31–48 (1973).

9. Young, H. M. et al. A single rostrocaudal colonization of the rodent intestine by enteric neuron precursors is revealed by the expression of Phox2b, Ret, and p75 and by explants grown under the kidney capsule or in organ culture. Dev. Biol. 202, 67–84 (1998).

10. Ganz, J. Gut feelings: Studying enteric nervous system development, function, and disease in the zebrafish model system. Dev. Dyn. Off. Publ. Am. Assoc. Anat. 247, 268–278 (2018).

11. Sasselli, V. et al. Planar cell polarity genes control the connectivity of enteric neurons. J. Clin. Invest. 123, 1763–1772 (2013).

12. Taraviras, S. & Pachnis, V. Development of the mammalian enteric nervous system. Curr. Opin. Genet. Dev. 9, 321–327 (1999).

13. Baynash, A. G. et al. Interaction of endothelin-3 with endothelin-B receptor is essential for development of epidermal melanocytes and enteric neurons. Cell 79, 1277–1285 (1994).

14. Zhu, Q. et al. Revealing a coherent cell state landscape across single cell datasets with CONCORD. BioRxiv Prepr. Serv. Biol. 2025.03.13.643146 (2025) doi:10.1101/2025.03.13.643146.

15. Lasrado, R. et al. Lineage-dependent spatial and functional organization of the mammalian enteric nervous system. Science 356, 722–726 (2017).

16. Lake, J. I. & Heuckeroth, R. O. Enteric nervous system development: migration, differentiation, and disease. Am. J. Physiol. Gastrointest. Liver Physiol. 305, G1–24 (2013).

17. Huycke, T. R. et al. Patterning and folding of intestinal villi by active mesenchymal dewetting. Cell 187, 3072–3089.e20 (2024).

18. Haber, A. L. et al. A single-cell survey of the small intestinal epithelium. Nature 551, 333–339 (2017).

19. McGinnis, C. S. et al. MULTI-seq: sample multiplexing for single-cell RNA sequencing using lipid-tagged indices. Nat. Methods 16, 619–626 (2019).

20. Wolock, S. L., Lopez, R. & Klein, A. M. Scrublet: Computational Identification of Cell Doublets in Single-Cell Transcriptomic Data. Cell Syst. 8, 281–291.e9 (2019).

21. Wolf, F. A., Angerer, P. & Theis, F. J. SCANPY: large-scale single-cell gene expression data analysis. Genome Biol. 19, 15 (2018).

22. Zhu, Q. et al. Revealing a coherent cell state landscape across single cell datasets with CONCORD. BioRxiv Prepr. Serv. Biol. 2025.03.13.643146 (2025) doi:10.1101/2025.03.13.643146.

23. Wolf, F. A., Angerer, P. & Theis, F. J. SCANPY: large-scale single-cell gene expression data analysis. Genome Biol. 19, 15 (2018).

24. Traag, V. A., Waltman, L. & van Eck, N. J. From Louvain to Leiden: guaranteeing well-connected communities. Sci. Rep. 9, 5233 (2019).

25. Smillie, C. S. et al. Intra- and Inter-cellular Rewiring of the Human Colon during Ulcerative Colitis. Cell 178, 714–730.e22 (2019).

26. Jin, S. et al. Inference and analysis of cell-cell communication using CellChat. Nat. Commun. 12, 1088 (2021).

27. Wolf, F. A., Angerer, P. & Theis, F. J. SCANPY: large-scale single-cell gene expression data analysis. Genome Biol. 19, 15 (2018).

28. Fang, Z., Liu, X. & Peltz, G. GSEApy: a comprehensive package for performing gene set enrichment analysis in Python. Bioinforma. Oxf. Engl. 39, btac757 (2023).

29. Kumar, N., Mishra, B., Athar, M. & Mukhtar, S. Inference of Gene Regulatory Network from Single-Cell Transcriptomic Data Using pySCENIC. Methods Mol. Biol. Clifton NJ 2328, 171–182 (2021).

30. Fattahi, F. et al. Deriving human ENS lineages for cell therapy and drug discovery in Hirschsprung disease. Nature 531, 105–109 (2016).

31. Barber, K., Studer, L. & Fattahi, F. Derivation of enteric neuron lineages from human pluripotent stem cells. Nat. Protoc. 14, 1261–1279 (2019).

32. Majd, H. et al. hPSC-Derived Enteric Ganglioids Model Human ENS Development and Function. 2022.01.04.474746 Preprint at 10.1101/2022.01.04.474746 (2022).

33. Dobin, A. et al. STAR: ultrafast universal RNA-seq aligner. Bioinforma. Oxf. Engl. 29, 15–21 (2013).

34. Liao, Y., Smyth, G. K. & Shi, W. featureCounts: an efficient general purpose program for assigning sequence reads to genomic features. Bioinforma. Oxf. Engl. 30, 923–930 (2014).

35. Love, M. I., Huber, W. & Anders, S. Moderated estimation of fold change and dispersion for RNA-seq data with DESeq2. Genome Biol. 15, 550 (2014).

36. Subramanian, A. et al. Gene set enrichment analysis: a knowledge-based approach for interpreting genome-wide expression profiles. Proc. Natl. Acad. Sci. U. S. A. 102, 15545–15550 (2005).

37. Korotkevich, G. et al. Fast gene set enrichment analysis. 060012 Preprint at 10.1101/060012 (2021).

38. Liberzon, A. et al. The Molecular Signatures Database (MSigDB) hallmark gene set collection. Cell Syst. 1, 417–425 (2015).

39. Benjamini, Y., Drai, D., Elmer, G., Kafkafi, N. & Golani, I. Controlling the false discovery rate in behavior genetics research. Behav. Brain Res. 125, 279–284 (2001).

40. Jin, S. et al. Inference and analysis of cell-cell communication using CellChat. Nat. Commun. 12, 1088 (2021).

41. Love, M. I., Huber, W. & Anders, S. Moderated estimation of fold change and dispersion for RNA-seq data with DESeq2. Genome Biol. 15, 550 (2014).

42. Shear, A. et al. Cerebral circulation improves with indirect bypass surgery combined with gene therapy. Brain Circ. 5, 119–123 (2019).

